# The dual nature of bacteriophage: growth-dependent predation and generalised transduction of antimicrobial resistance

**DOI:** 10.1101/2021.07.24.453184

**Authors:** Quentin J Leclerc, Jacob Wildfire, Arya Gupta, Jodi A Lindsay, Gwenan M Knight

## Abstract

Bacteriophage (“phage”) are both predators and evolutionary drivers for bacteria, notably contributing to the spread of antimicrobial resistance (AMR) genes by generalised transduction. Our current understanding of the dual nature of this relationship is limited. We used an interdisciplinary approach to quantify how these interacting dynamics can lead to the evolution of multi-drug resistant bacteria. We co-cultured two strains of Methicillin-resistant *Staphylococcus aureus*, each harbouring a different antibiotic resistance gene, with 80α generalized transducing phage. After a growth phase of 8h, bacteria and phage surprisingly coexisted at a stable equilibrium in our culture, the level of which was dependent on the starting concentration of phage. We detected double-resistant bacteria as early as 7h, indicating that transduction of AMR genes had occurred. We developed multiple mathematical models of the bacteria and phage relationship, and found that phage-bacteria dynamics were best captured by a model in which the phage burst size decreases as the bacteria population reaches stationary phase, and where phage predation is frequency-dependent. We estimated that one in every 10^8^ new phage generated was a transducing phage carrying an AMR gene, and that double-resistant bacteria were always predominantly generated by transduction rather than by growth. Our results suggest a shift in how we understand and model phage-bacteria dynamics. Although rates of generalised transduction could be interpreted as too rare to be significant, they are sufficient to consistently lead to the evolution of multi-drug resistant bacteria. Currently, the potential of phage to contribute to the growing burden of AMR is likely underestimated.

## Introduction

To counter the rapidly increasing global public health threat of antimicrobial resistance (AMR), we must urgently develop new solutions ^1^. “Phage therapy” is one such tool which has recently seen a renewed interest ^2^. This relies on using bacteriophage (or “phage”), major bacteria predators and the most abundant biological entities on the planet ^3^, as antimicrobial agents. However, phage are also natural drivers of bacterial evolution through horizontal gene transfer by “transduction” ^4,5^. AMR genes can be transferred by transduction at high rates, both *in vitro* and *in vivo* ^6–8^. The dual nature of phage (predation and transduction) makes them a double-edged sword in the fight against AMR, as they are themselves capable of contributing to the spread of the problem they aim to solve, yet our understanding of these dynamics and how to best represent them is limited.

There are two types of transduction; here, we focus on “generalised transduction”, which occurs during the phage lytic cycle, when non-phage genome DNA is mistakenly packaged in a new phage particle (Figure 1). The resulting transducing phage released upon lysis can then inject this genetic material into another bacterium. Current guidelines for phage therapy recommend that exclusively lytic phage should be used, removing the risk of the second type of transduction which relies on lysogeny (“specialised transduction”, where sections of bacterial DNA adjacent to the prophage integration site may be transferred upon excision of the prophage) ^9,10^. The possibility of generalised transduction remains, yet is currently often dismissed as too rare to be significant, despite being a common mechanism for the transfer of plasmids, major vectors of AMR genes ^4^. Alternatively, guidelines may state that the risk of generalised transduction should be minimised ^9,11,12^, yet there are currently no estimates or work quantifying what constitutes an “acceptable” risk. Previous reviews have highlighted the necessity to further investigate the potential impact of transduction in the context of phage therapy ^11–13^.

**Figure 1:**
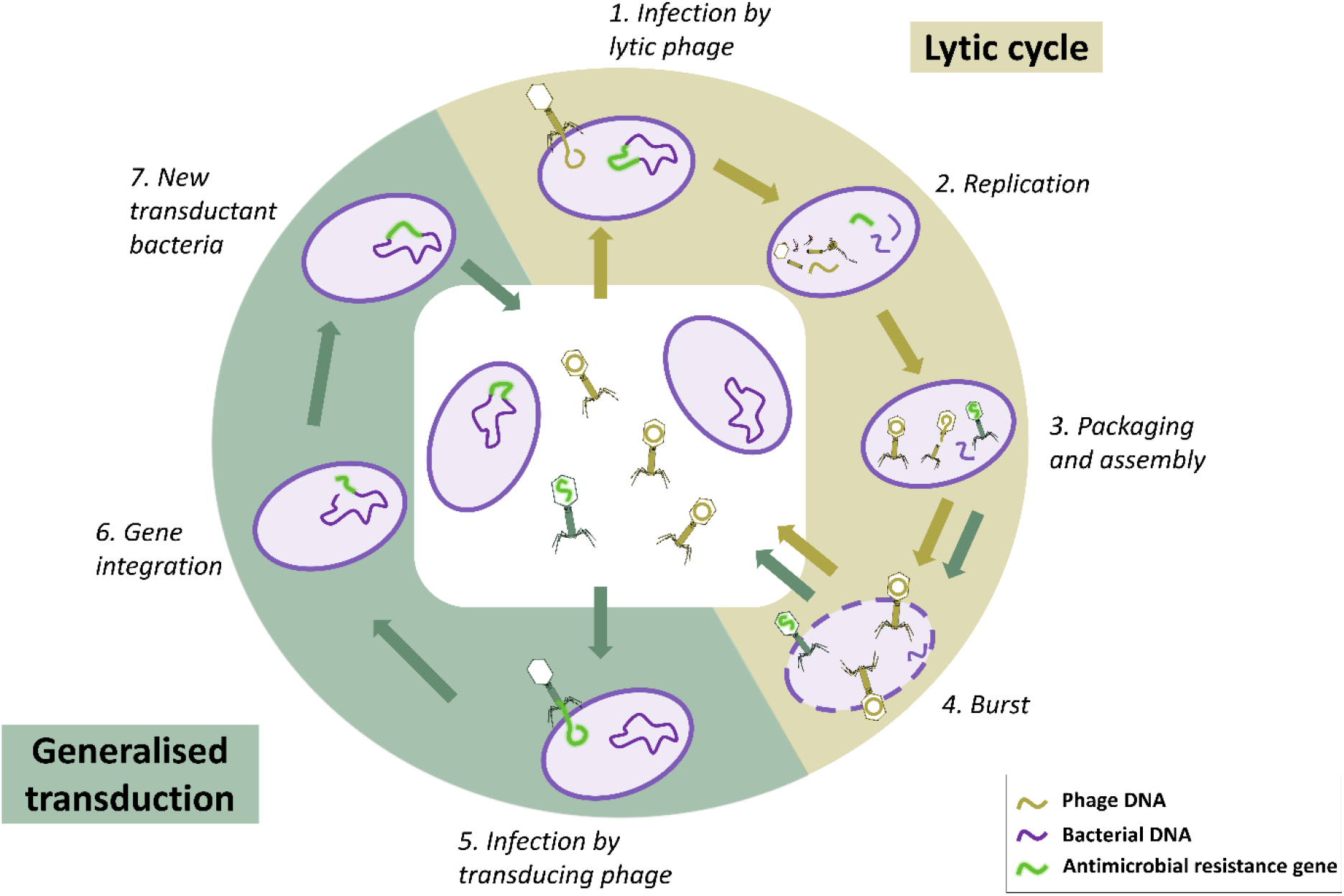
Phage lytic cycle and generalised transduction. In this environment, only some bacteria carry an antimicrobial resistance (AMR) gene (shown in green). The lytic cycle starts when a lytic phage infects a bacterium by binding and injecting its DNA (1). Phage molecules degrade bacterial DNA and utilise bacterial resources to create new phage components and replicate (2). These components are then assembled to form new phage particles (3). At this stage, bacterial DNA leftover in the cell can be packaged by mistake instead of phage DNA, which creates a transducing phage and starts the process of generalised transduction. In our example, the transducing phage carries the AMR gene. After a latent period of typically several minutes, the phage trigger lysis of the bacterium, bursting it and releasing the phage (4). The transducing phage can infect another bacterium, binding and injecting the AMR gene it is carrying (5). If this gene is successfully integrated into the bacterial chromosome (6), this creates a new transductant bacterium carrying this AMR gene (7). Note that the transduced bacterial DNA could also be a plasmid, in which case it would circularise instead of integrating into the chromosome of the transductant bacterium. Not to scale.

Mathematical models have been used to gain insights into phage predation dynamics which cannot be obtained solely with experimental work, such as rates of predation and optimal conditions for phage to clear bacteria ^14^. Such models typically assume a density-dependent interaction, with new phage infections calculated as the number of susceptible bacteria, multiplied by the number of phage and an adsorption constant ^14–16^. This approach has limitations, as density-dependent models have failed to predict equilibriums observed in some *in vitro* conditions between phage and *E. coli* ^17^. Moreover, phage and bacterial replication are likely to be linked, as they both rely on the bacterial machinery; phage predation may slow as bacteria reach stationary phase ^14,17–23^. However, this is a feature which is not commonly included in mathematical models of phage-bacteria dynamics ^14^. Finally, models often only rely on data of phage-bacteria interactions measured once per day, or for a few hours ^17–19,24^. A current lack of detailed data means that capturing these underlying dynamics which occur in less than an hour has not yet been possible.

To the best of our knowledge, only three modelling studies have included transduction of AMR genes ^25–27^. All three modelled complex environments, including resistance to phage, antibiotics, and both lytic and lysogenic cycles. This complexity, combined with the fact that these studies were not paired with complementary *in vitro* or *in vivo* data, means that they relied on assumptions and previously published estimates, instead of parameter values derived from a single environment and set of conditions. This limits the wider reliability of conclusions made using these models ^13^.

In this article, we investigate the dual nature of phage dynamics using the clinically relevant bacteria Methicillin-resistant *Staphylococcus aureus* (MRSA) ^28^. Transduction is the main mechanism of horizontal gene transfer driving evolution for these bacteria ^29^, and phage therapy is currently being investigated to treat MRSA infections ^30,31^. We aim to clarify the dual nature (predation and transduction) of the interaction between MRSA and phage by generating novel *in vitro* data, identifying biologically plausible hypotheses which may explain the dynamics seen, and developing mathematical models to test the validity of these hypotheses in our system.

## Results

### Transduction and phage predation dynamics *in vitro*

We focused on two laboratory strains of *Staphylococcus aureus*, each resistant to either erythromycin (and referred to as B_E_) or tetracycline (B_T_). In our experimental conditions, the antimicrobial resistance (AMR) genes can only be transferred between bacteria by generalised transduction mediated by exogenous phage. Transduction of either AMR gene to the other strain will result in the formation of double-resistant progeny (referred to as B_ET_).

We conducted a co-culture with only the two single-resistant strains and exogenous temperate phage 80α (P_L_) capable of generalised transduction. We grew the bacteria and phage over 24h, with hourly counts of bacteria and lytic phage between 0-8h and 16-24h. The starting concentration of bacteria was 10^4^ colony-forming units (cfu) per mL, and of phage was approximately either 10^3^, 10^4^ or 10^5^ plaque-forming units (pfu) per mL, equivalent to multiplicities of infection of 0.1, 1 and 10 (defined as starting ratio of phage to bacteria ^32^).

We detected double-resistant progeny (B_ET_) as early as 7h in our co-cultures, indicating that transfer of AMR genes by generalised transduction had occurred (Figure 2). B_ET_ numbers remained below 100 cfu/mL after 24h, but were consistently generated in each of our experimental replicates. Colonies of double-resistant progeny were screened using polymerase chain reaction (PCR) to confirm that they contained both resistance genes *ermB* and *tetK*, and had not instead gained resistance to either antibiotic by mutation (Figure 2 – figure supplement 1).

**Figure 2:**
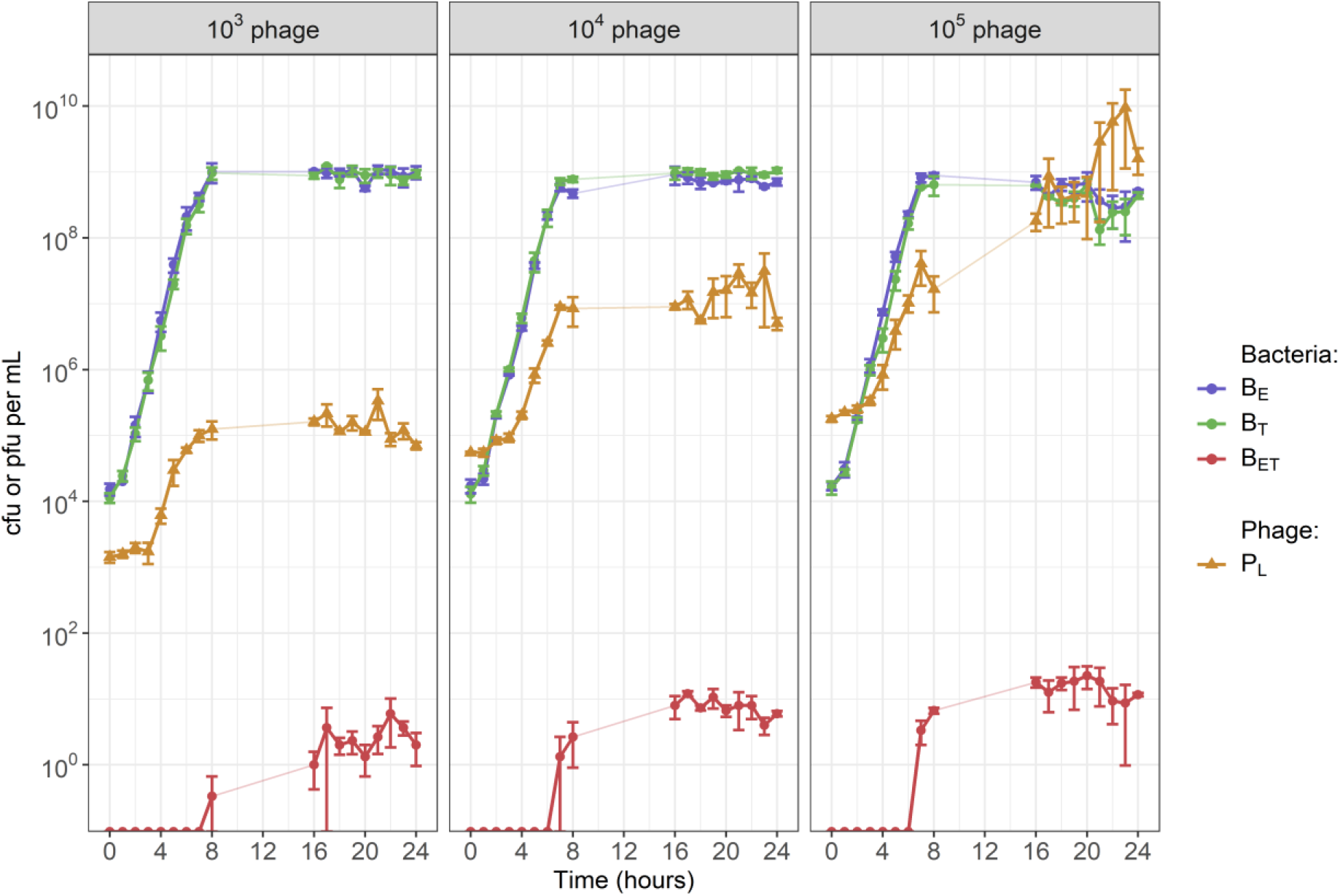
The starting concentration of exogenous phage 80α affected the equilibrium values of phage and bacteria in our co-cultures. The starting concentration of both single-resistant *S. aureus* parent strains (B_E_ to erythromycin & B_T_ to tetracycline) was 10^4^ colony-forming units (cfu) per mL. Each panel shows the results with a different starting concentration of exogenous phage 80α (P_L_): either 10^3^, 10^4^ or 10^5^ plaque-forming units (pfu) per mL. We detected double-resistant progeny (B_ET_) as early as 7h, indicating that transduction occurred rapidly. Error bars indicate mean +/− standard error, from 3 experimental replicates. There is no data for the time period 9h-15h. **Figure supplement 1: Confirmation of double-resistant progeny by polymerase-chain reaction.** Five single colonies were sampled from a double antibiotic plate (1-5), containing bacteria plated after 24h of co-culture started only with single-resistant parent strains (E and T) and exogenous phage. L: ladder; E: erythromycin resistance gene (*ermB*); T: tetracycline resistance gene (*tetK*). **Figure supplement 2: Growth curves for bacteria in the absence of exogenous phage.** Solid lines correspond to *in vitro* data, and dashed lines to the model output generated using the median values of the parameter distributions obtained by model fitting. Shaded areas indicate error obtained by resampling the model results from a Poisson distribution ten times. **Figure supplement 3: Transduction co-culture datasets overlaid.**

The starting concentration of exogenous phage affected whether phage and bacteria were able to reach an equilibrium and co-exist without increasing or decreasing in our co-cultures (Figure 2). With a starting concentration of either 10^3^ or 10^4^ pfu/mL, lytic phage reached a steady-state after 8h (at approximately 10^5^ pfu/mL for a starting concentration of 10^3^, and 10^7^ pfu/mL for 10^4^). In both cases, bacteria replicated for 8h before reaching a steady-state around 10^9^ cfu/mL, similar to what was seen in the absence of exogenous phage (Figure 2 – figure supplement 2). With a starting phage concentration of 10^5^ pfu/mL, we did not see an equilibrium between phage and bacteria, as phage numbers kept increasing up to 10^10^ pfu/mL by 24h, and bacteria numbers started decreasing after 20h. The datasets are shown overlaid in Figure 2 – figure supplement 3.

### Absence of lysogeny in our co-culture

80α is a temperate phage, which means that it may undergo lysogeny and integrate in the bacterial chromosome as a prophage ^33^. This would grant lysogenic immunity to the bacteria, preventing further lytic infection by 80α, and potentially explaining why bacteria and phage densities reached steady-states in our experiments (Figure 2). To investigate this, we initiated co-cultures with 10^4^ cfu/mL of each single-resistant strain and 10^4^ pfu/mL of 80α phage. After 24h, we diluted these in fresh media to form new co-cultures with an inocula of 10^4^ cfu/mL of bacteria already exposed to the phage, and added phage to reach a concentration of 10^4^ pfu/mL again. We did not see any difference in phage and bacteria numbers after 24h regardless of whether or not the bacteria had been previously exposed to phage (Figure 3a).

**Figure 3:**
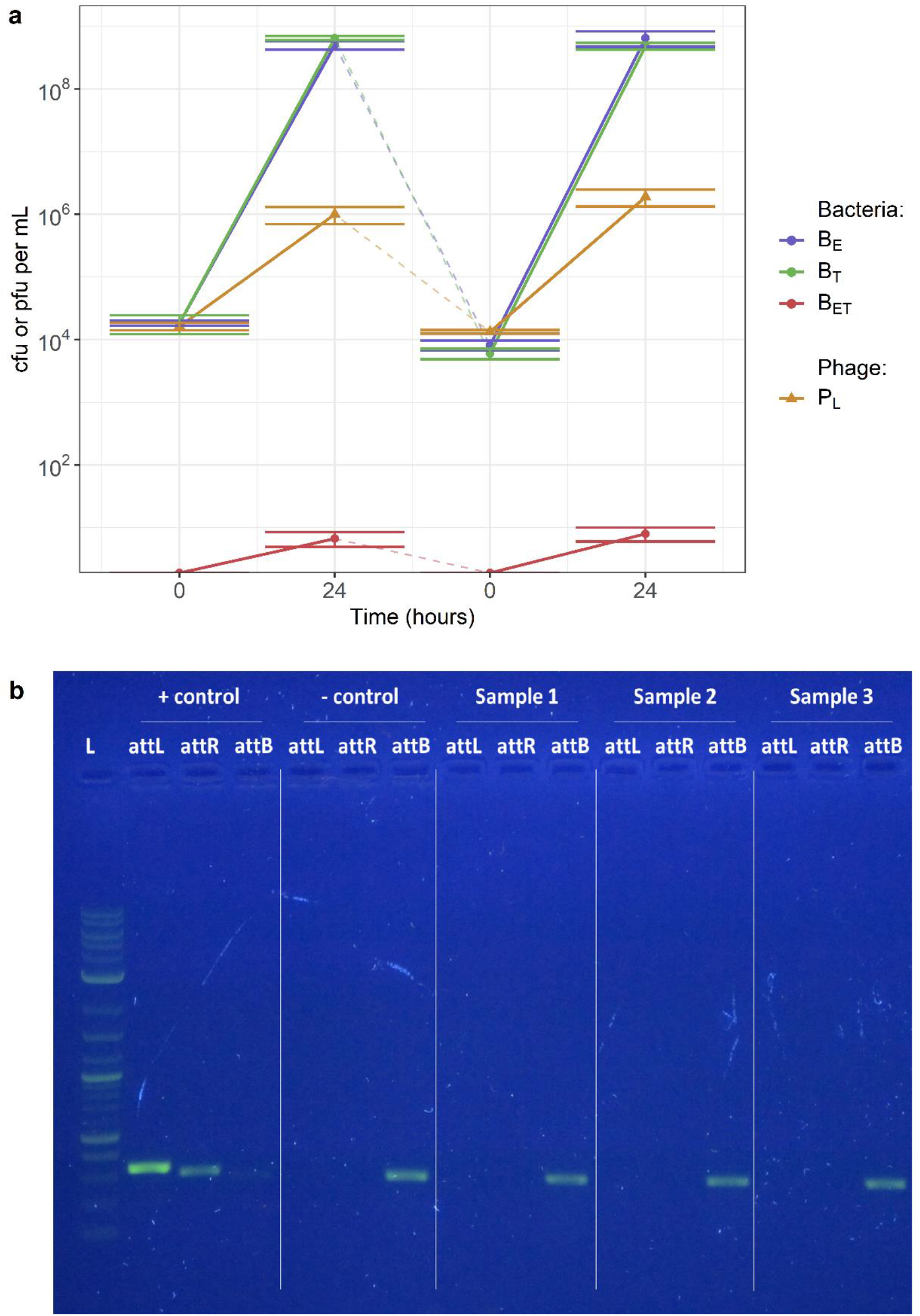
80α lysogeny does not occur at a detectable level in our co-culture. (a) Co-cultures with bacteria not exposed or previously exposed to phage. The starting concentration of both single-resistant *S. aureus* parent strains (B_E_ to erythromycin & B_T_ to tetracycline) was 10^4^ colony-forming units (cfu) per mL, and the starting concentration of exogenous phage 80α (P_L_) was 10^4^ plaque-forming units (pfu) per mL. Double-resistant progeny (B_ET_) are generated by transduction. The initial co-culture was diluted in fresh media after 24h, to form a new co-culture with bacteria previously exposed to phage. Phage were added in the new co-culture to reach a concentration of 10^4^ pfu/mL. Error bars indicate mean +/− standard error, from 3 experimental replicates. **(b) Confirmation of absence of detectable lysogeny by polymerase-chain reaction.** DNA was extracted from the co-cultures after 24h. *S. aureus* RN4220 strains lysogenic and non-lysogenic for 80α were used as positive and negative controls. L: ladder; *attL*: left prophage junction; *attR*: right prophage junction; *attB*: bacterial insertion site. Detection of *attL* and *attR* indicates that prophage are present in the DNA, while detection of *attB* indicates the presence of bacteria not lysogenic for 80α

In addition, we extracted DNA from the co-cultures after 24h, and conducted PCRs targeting the prophage junctions (*attL* and *attR*) and bacterial insertion site (*attB*). Only the intact bacterial insertion site was detected in our samples, indicating an absence of prophage in our bacteria (Figure 3b). Overall, these results suggest that lysogeny does not occur at a detectable level in our co-culture, and that we can exclude any dynamics relating to lysogeny in our analysis and model below.

### Bacterial growth estimates in the absence of exogenous phage

When grown together in the absence of exogenous phage, single and double resistant bacteria replicated exponentially and reached stationary phase after 8h at 10^9^ colony-forming units (cfu) per mL (Figure 2 – figure supplement 2). For each pair of strains, we estimated relative fitness *W* using Equation 1.

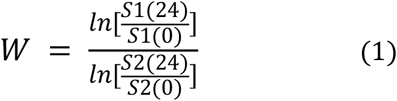

Where S1(*t*) and S2(*t*) represent the number of bacteria (in cfu/mL) from the chosen strains 1 and 2, at times *t* = 0 or 24 hours.

B_E_ did not show a significant fitness cost relative to B_T_ over 24h of growth (mean relative fitness 1.02, sd 0.03). The double-resistant progeny B_ET_ did not show a significant fitness cost relative to either single-resistant parent strain (mean relative fitness to B_E_: 0.96, sd 0.06; mean relative fitness to B_T_: 0.98, sd 0.03).

We obtained maximum growth rate estimates by fitting a logistic growth model to the *in vitro* data. This is shown in Equation 2, where μ_max_ is the maximum bacterial growth rate, B_max_ is the carrying capacity, and *B* the concentration of bacteria (in cfu/mL).

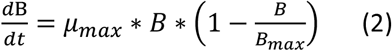

The median estimated maximum growth rates were 1.61 for B_E_ (95% credible interval 1.59-1.63), 1.51 for B_T_ (1.49-1.53) and 1.44 for B_ET_ (1.42-1.47), with a total carrying capacity of 2.76 × 10^9^ cfu/mL (2.61 × 10^9^ – 2.98 × 10^9^).

### Investigation of possible phage-bacteria interactions using a flexible modelling framework

#### Model structure

We designed a mathematical model to reproduce the *in vitro* phage-bacteria dynamics, including generalised transduction of resistance genes. During our experiment, our co-culture contained up to three strains of bacteria: the two single-resistant parents (B_E_, B_T_) and the double-resistant progeny (B_ET_). Although we were only able to count lytic phage (P_L_), based on the biology of generalised transduction (Figure 1) we know that there were also transducing phage carrying either the erythromycin resistance gene (P_E_), or the tetracycline resistance gene (P_T_). Since we did not detect any evidence of 80α lysogeny in our co-culture after 24h, we did not include this feature in the model. The corresponding model diagram is shown in Figure 4a. The complete model equations can be found in Methods.

**Figure 4:**
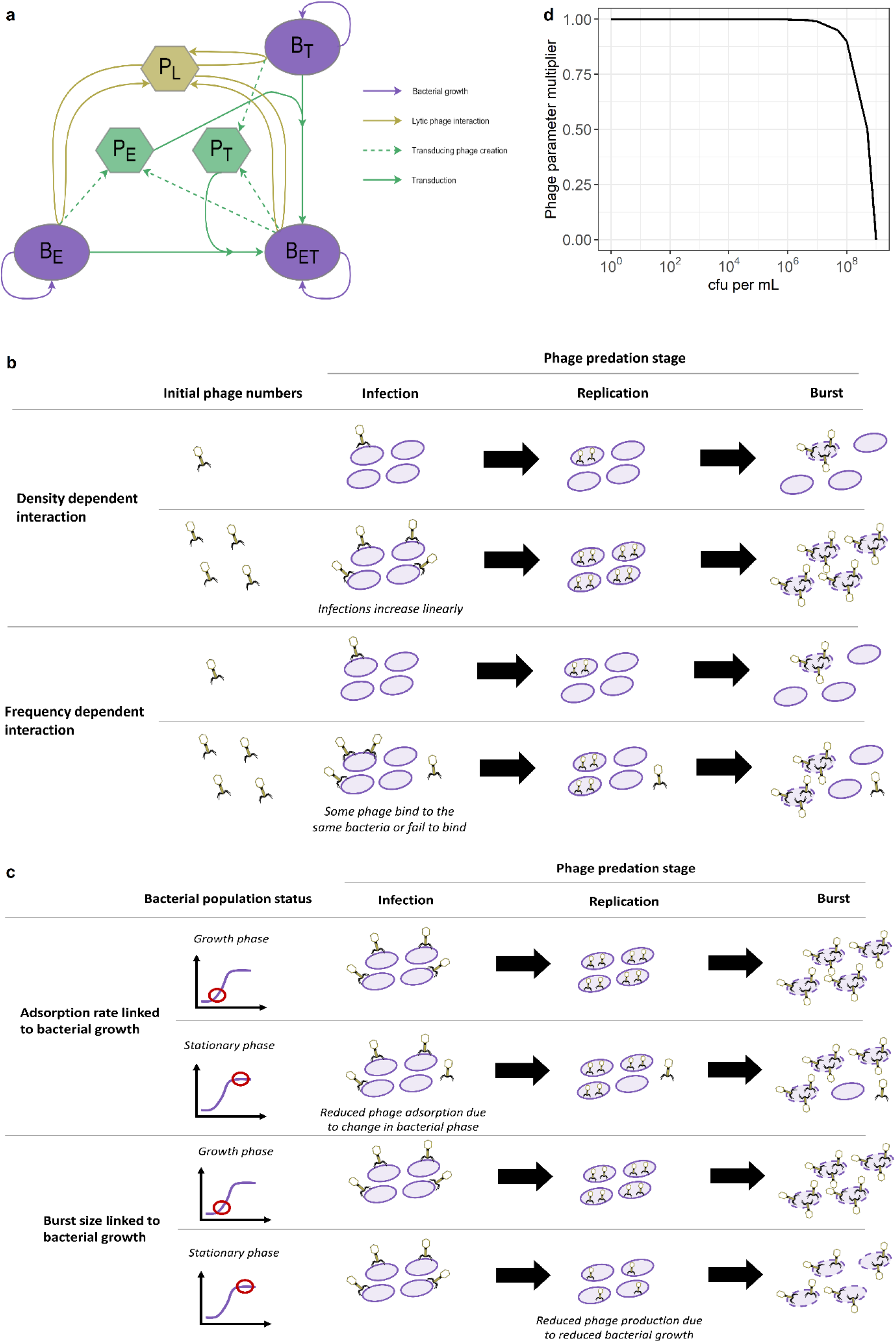
Phage predation and generalised transduction model diagram, and different phage-bacteria interactions considered. (**a) Model diagram.** Each bacteria strain (B_E_ resistant to erythromycin, B_T_ resistant to tetracycline, or B_ET_ resistant to both) can replicate (purple). The lytic phage (P_L_) multiply by infecting a bacterium and bursting it to release new phage (gold). This process can create transducing phage (P_E_ or P_T_) carrying a resistance gene (*ermB* or *tetK* respectively) taken from the infected bacterium (green). These transducing phage can then generate new double resistant progeny (B_ET_) by infecting the bacteria strain carrying the other resistance gene (green). **(b) Phage predation in the model is either density- or frequency-dependent.** With a density-dependent interaction, the number of infections scales linearly with the number of phage and bacteria (top). A frequency-dependent interaction illustrates that some phage may not infect a bacterium, or that multiple phage may infect the same bacterium (bottom). **(c) Phage predation in the model can decrease as bacterial growth decreases.** A change in bacterial growth phase can affect surface receptors, leading to a reduced phage adsorption rate (top). Since phage replication relies on bacterial processes, a reduced bacterial growth can translate into a reduced phage burst size (bottom). **(d) Proposed function linking phage predation parameters to bacterial growth.** This shows the multiplier applied to decrease phage parameters as the bacterial population increases towards carrying capacity, equivalent to a decrease in bacterial growth. Here, the carrying capacity is 2.76 × 10^9^ colony-forming units (cfu)/mL, estimated from our data.

Using this modelling framework, we explored a combination of different phage-bacteria interactions, described below (Figure 4b-c). By fitting the models to our experimental data, we could rule out certain interactions and suggest the best model to reproduce the phage-bacteria dynamics seen *in vitro*.

#### First phage-bacteria interaction: density versus frequency-dependent phage predation

The most common approach to model phage-bacteria dynamics is to assume that phage predation is density-dependent ^14^. This means that, over one time step, the number of phage infecting bacteria and the number of bacteria infected by phage are both equal to the product of the number of bacteria (B), phage (P), and phage adsorption rate (β), as shown in equation (3).

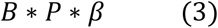

The density-dependent interaction implies that the number of new infections scales linearly with the number of phage and bacteria (Figure 4b). Therefore, even if we keep a constant number of phage, increasing bacteria numbers always leads to a linear increase in the estimated number of new infections. Although this simplification is useful and holds for a range of values, it has been suggested that it is not biologically realistic for small numbers of phage or bacteria, since in reality one phage can only infect one bacterium over one time step ^17^.

To overcome these issues, we consider an alternative interaction, where phage predation is frequency-dependent ^34^. This accounts for the fact that one phage does not necessarily always lead to one infection. For example, phage may sometimes fail to bind to bacteria, or multiple phage may bind to the same bacterium ^32^ (Figure 4b). Importantly, this mathematical interaction guarantees that, at any given time point, the number of phage infecting bacteria and the number of bacteria infected by phage can never be greater than the total number of phage or bacteria in the system. Over one time step, the proportion of phage infecting any bacteria (λ) is defined by equation (4).

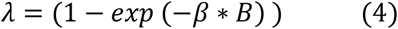

Similarly, the proportion of bacteria being infected by at least one phage (φ) is calculated with equation (5).

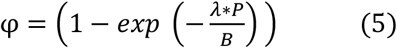

#### Equilibrium analyses for the density and frequency-dependent models

Despite these being common methods to represent phage-bacteria interactions in mathematical models, previous analyses have suggested that the density and frequency-dependent interactions alone cannot capture the equilibrium levels we and others have seen ^18,35^. We explore this in the context of our own *in vitro* data using equilibrium analyses.

Assuming that transduction and the phage latent period are negligible, a simplified model representing phage predation as a density-dependent process is shown in equations (6) and (7).

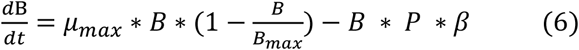

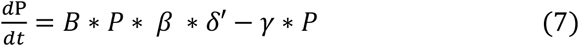

Where μ_max_ is the maximum bacterial growth rate, B_max_ is the carrying capacity, β is the phage adsorption rate, γ is the phage decay rate, and δ is the phage burst size, with δ’ equal to δ − 1. To solve for equilibrium (i.e. 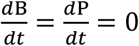), equations (6) and (7) can be rewritten as equations (8) and (9).

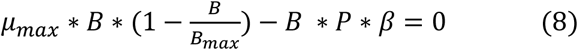

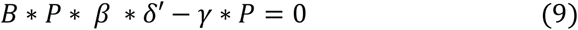

Since we are interested in an equilibrium with the condition that there are still bacteria and phage in the environment (i.e. B≠0 and P≠0), we can divide equations (8) and (9) by B and P respectively to obtain equations (10) and (11). These must hold true for there to be a non-zero bacteria and phage population at equilibrium.

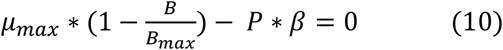

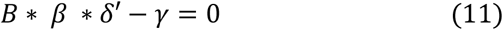

We then obtain equations (12) and (13) by rearranging (10) and (11) to give expressions for P and B at equilibrium.

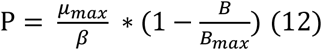

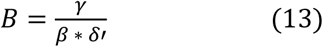

In our experiment with a starting phage concentration of 10^4^ plaque-forming units (pfu)/mL, after 24h the bacteria concentration was approximately 10^9^ colony-forming units (cfu)/mL, and the phage concentration was 10^5^ pfu/mL. If we replace the corresponding terms in equations (12) and (13) with these values, alongside the carrying capacity (2.8 × 10^9^) and average of the growth rates estimated (1.52), we obtain equations (14) and (15).

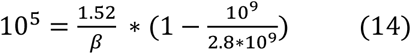

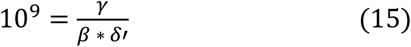

Rearrangement of equation (14) leads to a solution for phage adsorption (β) (equation (16)).

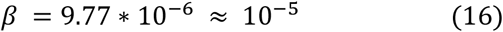

Substituting this into equation (15) leads to a value for phage decay rate (γ) (equation (17)).

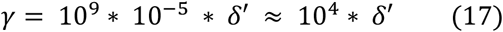

Giving rise to the condition that the phage decay rate γ must be approximately 10^4^ times greater than the burst size δ’. As at least one phage must be released upon bursting, the burst size δ’ is greater than 1. However, the phage decay rate γ, which represents the proportion of phage inactivated during one time-step, must be less than 1 and hence this is impossible.

As for the frequency-dependent interaction, a simplified model of this process is shown in equations (18) and (19).

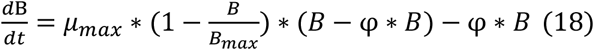

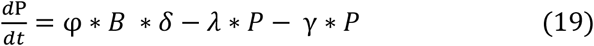

With the condition that 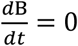 equation (18) can be rewritten as equation (20). This condition must hold in order for there to be a non-zero equilibrium.

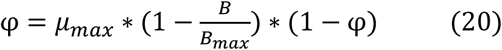

Using our equilibrium values (B = 10^9^, P = 10^5^, B_max_ = 2.8 × 10^9^, μ_max_ = 1.52), this is equivalent to equation (21).

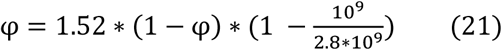

Equation (21) is solved to obtain a value for φ (equations (22) – (25)).

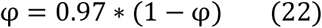

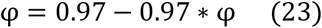

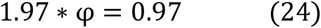

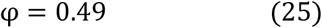

Using our equilibrium values and this value of 0.49 for φ, equation (5) can be rewritten as equation (26).

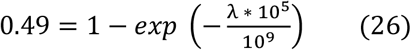

Equation (26) is solved to obtain a value for λ (equations (27) – (31)).

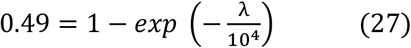

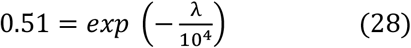

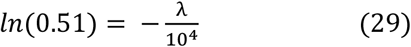

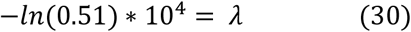

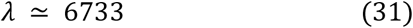

Therefore, λ must be approximately equal to 6733 to obtain a non-zero equilibrium as seen in our *in vitro* data with our model fitted values (biologically plausible ones). However, the definition of λ according to equation (4) implies that λ < 1. Therefore, this is impossible.

Even though these analyses rely on a simplified set of equations, using realistic parameter values we have shown that a non-zero equilibrium, as we have seen *in vitro,* cannot be reproduced using models with only a density or frequency-dependent interaction. Instead, phage-bacteria co-existence may be explained by variations in phage predation parameters depending on bacterial resources availability, or bacterial growth rate ^14,17–22^. However, to the best of our knowledge a simple mathematical expression linking phage predation to bacterial growth has not yet been developed.

#### Second phage-bacteria interaction: dependence of phage predation on bacterial growth

Here, we consider that a decrease in bacterial growth as bacteria reach stationary phase could firstly affect the phage adsorption rate β, due to changes in receptors on bacterial surfaces, which affect opportunities for phage to bind (Figure 4c). Secondly, this could affect phage production, and thus burst size δ, as phage replication relies on bacterial processes and may decrease when bacterial growth slows down (Figure 4c). Using a single phage predation multiplier, with the same principle of logistic growth applied to bacteria, we allow either or both β and δ to decrease as bacterial growth decreases in our model (equations (32) and (33)).

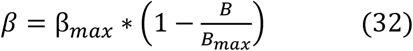

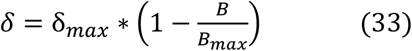

These equations imply that, as bacterial population size increases towards carrying capacity (B_max_), phage parameters will be reduced (Figure 4d).

### Identification of the best-fitting phage-bacteria interactions to reproduce the *in vitro* dynamics

Overall, we considered 6 different models, either density- or frequency-dependent, and with either or both the phage adsorption rate and burst size linked to bacterial growth. Note that we did not include a phage decay rate in these models, as this did not affect the dynamics of the system over 24h, for a wide range of decay rates (Figure 5 – figure supplement 1).

We used a Bayesian methodology to fit the models simultaneously to the lytic phage and double-resistant progeny numbers from the transduction co-culture datasets with starting phage concentrations of 10^3^ and 10^5^ pfu/mL (Figure 2), and tested whether the estimated parameters could reproduce the dynamics seen with the starting phage concentration of 10^4^ pfu/mL. Convergence and posterior distribution plots for our best-fitting model are shown in Figure 5 – figure supplement 2.

All models successfully reproduced the trends seen *in vitro* when the phage were started at either 10^3^ and 10^4^ pfu/mL (Figure 5a-b). However, only the two models where only phage burst size decreases as the bacteria population approaches carrying capacity were able to reproduce the increase in phage numbers seen in the later hours of the 10^5^ pfu/mL dataset, despite all models having been fitted to this dataset (Figure 5a-b). This was confirmed by calculating the average Deviance Information Criteria (DIC) value for the models, which favours best-fitting models while penalising more complex models (i.e. with more parameters) ^36^. The two models where only phage burst size decreases as the bacteria population approaches carrying capacity had the lowest DIC values, indicating that they were the better-fitting models (Table 1).

**Figure 5:**
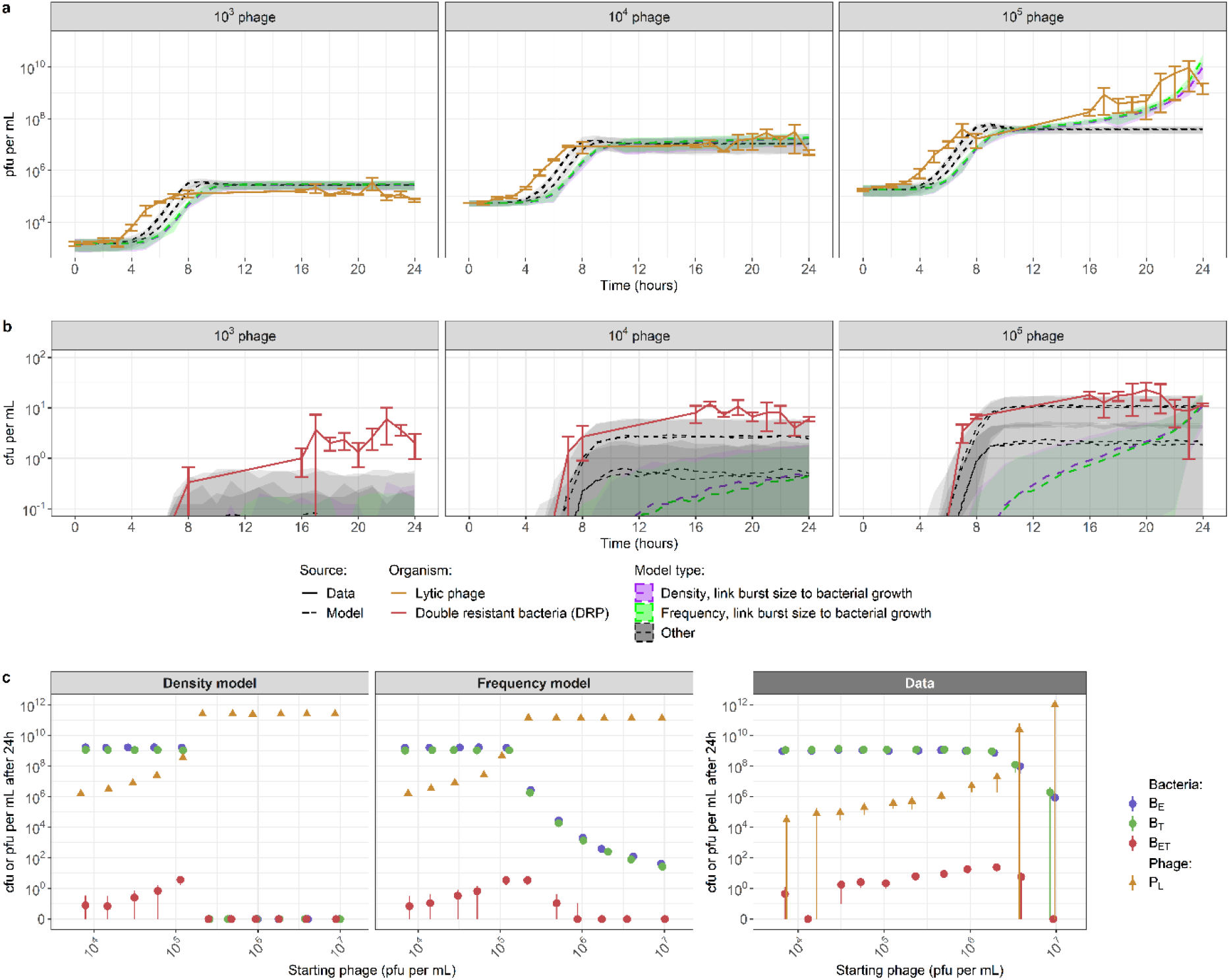
Accuracy of the best-fitted models to reproduce *in vitro* phage-bacteria dynamics. (a-b) The models with only phage burst size linked to bacterial growth are the most accurate to reproduce *in vitro* trends in lytic phage (a) and double resistant bacteria (b) numbers, starting from a bacteria concentration of 10^4^ cfu/mL and varying phage concentrations. All models (dashed lines) can reproduce the trends seen *in vitro* when phage are started at 10^3^ or 10^4^ pfu/mL (data in solid lines), but only the models with just the phage burst size linked to bacterial growth (coloured model output) can reproduce the trend seen when phage are started at 10^5^ pfu/mL. Other models (grey) either only have the phage adsorption rate linked to bacterial growth, or both the phage adsorption rate and burst size. Models are fitted to the 10^3^ and 10^5^ data, and tested with the 10^4^ data. Parameter values used are the median fitted values (Table 1). Shaded areas indicate standard deviation generated from Poisson resampling of model results. Error bars for the data (solid lines) indicate mean +/− standard error, from 3 experimental replicates. **(c) When further testing fitted model dynamics starting from 10^6^ cfu/mL bacteria and varying phage concentrations, the density-dependent model incorrectly predicts bacterial extinction, while the frequency-dependent model reproduces the trend, but not the exact values of the 24h data.** In the co-culture used to generate the data, each single-resistant parent strain (B_E_ and B_T_) is added at a starting concentration of 10^6^ cfu/mL, and no double-resistant progeny (B_ET_) are initially present. The starting concentration of lytic phage (P_L_) varies (x axis). Points indicate mean results, and are each slightly shifted horizontally to facilitate viewing. Error bars indicate either mean +/− standard deviation for the models (left/centre panels), or mean +/− standard error for the data (right panel). Parameter values used are the median fitted values (Table 1). **Figure supplement 1: Model results are not affected by phage decay rate over a wide range of values.** Previous estimates of phage decay rate per hour are between 10^−3^ *in vitro* and up to 10^−1^ *in vivo* ^38^. **Figure supplement 2: Convergence and posterior distributions for the best-fitting model.** (a-d) Parameter convergence plots. Fitting was performed using two chains (black and red). (e-h) Posterior distributions. The prior distributions for the burst size and latent period are shown in blue. The prior distributions for other parameters were uninformative and not shown. **Figure supplement 3: Model performance in reproducing the 24h data values for different starting concentrations of phage with different links between phage predation and bacterial growth rate.** (a) Phage adsorption rate decreases as bacterial growth rate decreases. (b) Phage burst size and adsorption rate decrease as bacterial growth rate decreases.

**Table 1:** Estimated parameter values from fitting to *in vitro* data. Values show median and 95% credible intervals for posterior distributions. Parameter units are indicated in parentheses. Fitting was performed using the Markov chain Monte Carlo Metropolis–Hastings algorithm and the data from the co-culture with a starting bacterial concentration of 10^4^ cfu/ml and phage concentration of 10^3^ and 10^5^ pfu/ml. DIC: Deviance Information Criteria. A smaller DIC indicates better model fit. DIC values are relative to the smallest DIC calculated, which is for the frequency-dependent model with only burst size linked to bacterial growth (line 5, parameters in bold).

| Interaction type | Adsorption rate linked to growth | Burst size linked to growth | Adsorption rate $\beta$ (phage<sup>-1</sup> bacteria<sup>-1</sup> hour<sup>-1</sup>) | Burst size $\delta$ (phage) | Transducing phage proportion $\alpha$ (proportion of burst size) | Phage latent period $\tau$ (hour) | DIC |
| --- | --- | --- | --- | --- | --- | --- | --- |
| Density dependent | Yes | No | $4.5 \times 10^{-9}$ ($4.1 \times 10^{-9}$ ; $5.0 \times 10^{-9}$) | 12 (10 ; 14) | $3.1 \times 10^{-8}$ ($1.5 \times 10^{-8}$ ; $5.8 \times 10^{-8}$) | 0.64 (0.55 ; 0.73) | 610 |
| | No | Yes | $1.6 \times 10^{-10}$ ($1.5 \times 10^{-10}$ ; $1.7 \times 10^{-10}$) | 79 (72 ; 86) | $1.4 \times 10^{-8}$ ($1.1 \times 10^{-8}$ ; $1.7 \times 10^{-8}$) | 0.65 (0.62 ; 0.69) | 63 |
| | Yes | Yes | $4.3 \times 10^{-9}$ ($3.9 \times 10^{-9}$ ; $4.6 \times 10^{-9}$) | 43 (37 ; 49) | $1.2 \times 10^{-8}$ ($6.4 \times 10^{-9}$ ; $2.3 \times 10^{-8}$) | 0.93 (0.86 ; 0.99) | 298 |
| Frequency dependent | Yes | No | $5.1 \times 10^{-9}$ ($3.7 \times 10^{-9}$ ; $6.7 \times 10^{-9}$) | 10 (8 ; 12) | $3.1 \times 10^{-7}$ ($2.3 \times 10^{-7}$ ; $4.3 \times 10^{-7}$) | 0.60 (0.50 ; 0.69) | 680 |
|  | No | Yes | **$2.3 \times 10^{-10}$ ($2.1 \times 10^{-10}$ ; $2.4 \times 10^{-10}$)** | **76 (70 ; 83)** | **$1.0 \times 10^{-8}$ ($8.5 \times 10^{-9}$ ; $1.4 \times 10^{-8}$)** | **0.72 (0.69 ; 0.77)** | **0** |
| | Yes | Yes | $4.7 \times 10^{-9}$ ($3.8 \times 10^{-9}$ ; $5.8 \times 10^{-9}$) | 31 (26 ; 37) | $1.7 \times 10^{-7}$ ($1.3 \times 10^{-7}$ ; $2.1 \times 10^{-7}$) | 0.88 (0.79 ; 0.96) | 370 |

Our initial experiments considered the dynamics over 24h for varying phage starting concentrations. To test the ability of our model to recreate the dynamics under changing bacterial levels, we replicated our transduction co-culture experiments with starting concentrations of 10^6^ cfu/mL bacteria instead of 10^4^ cfu/mL, varying the starting phage concentration between 10^4^ and 10^6^ pfu/mL, and measuring bacteria and phage numbers after 24h of co-culture. We then used the estimated parameter values (Table 1) to try to reproduce these 24h numbers of bacteria and phage.

Increasing the starting phage concentration led to an increase in the number of phage after 24h (Figure 5c). For a starting phage concentration between 10^4^ and 10^6^ pfu/mL, increasing starting phage numbers did not affect single-resistant parents B_E_ and B_T_ numbers after 24h, but led to a progressive increase in double-resistant progeny B_ET_ numbers. Increasing starting phage numbers above 10^6^ pfu/mL caused bacteria numbers after 24h to decrease.

Using the estimated parameter values (Table 1) with the model where only burst size is linked to bacterial growth, we see that the density model cannot reproduce these dynamics as it predicts that bacteria become extinct rapidly (Figure 5c). The frequency-dependent model is able to reproduce these trends, but fails to recreate the exact same numbers of phage and bacteria, predicting a decline in bacterial levels when the starting phage concentration increases above 10^5^ pfu/mL, a lower threshold than seen in the data (Figure 5c). The same overall trends are seen for the models where only the adsorption rate is linked to bacterial growth, or both adsorption rate and burst size (Figure 5 – figure supplement 3).

### Analysis of phage predation and transduction dynamics

Parameter estimates for our best-fitting model (with a frequency-dependent interaction and a link between phage burst size and bacterial growth only) suggest that the adsorption rate is 2.3 × 10^−10^ (95% credible interval: 2.1 × 10^−10^ – 2.4 × 10^−10^) which is the smallest estimate from the models (Table 1). On the other hand, the estimated burst size is relatively large at 76 (70 – 83) phage, and is higher than a previous *in vitro* estimate for 80α of 40 ^37^. However, due to the decrease in burst size when bacteria are in stationary phase, we expect that this number would change depending on the conditions under which it is measured (Figure 6a). Finally, the estimated latent period of 0.72h (0.69 – 0.77) is slightly longer than a previous in vitro estimate of 0.67h ^37^. Regarding the other models, we note some biologically unlikely parameter estimates which further suggest that these models are inappropriate, such as the low burst size for the models with only the adsorption rate linked to bacterial growth (12 (10 – 14) and 10 (8 – 12)), or the high latent period for the models with both adsorption rate and burst size linked to bacterial growth (0.93 (0.86 – 0.99) and 0.88 (0.79 – 0.96)) (Table 1).

**Figure 6:**
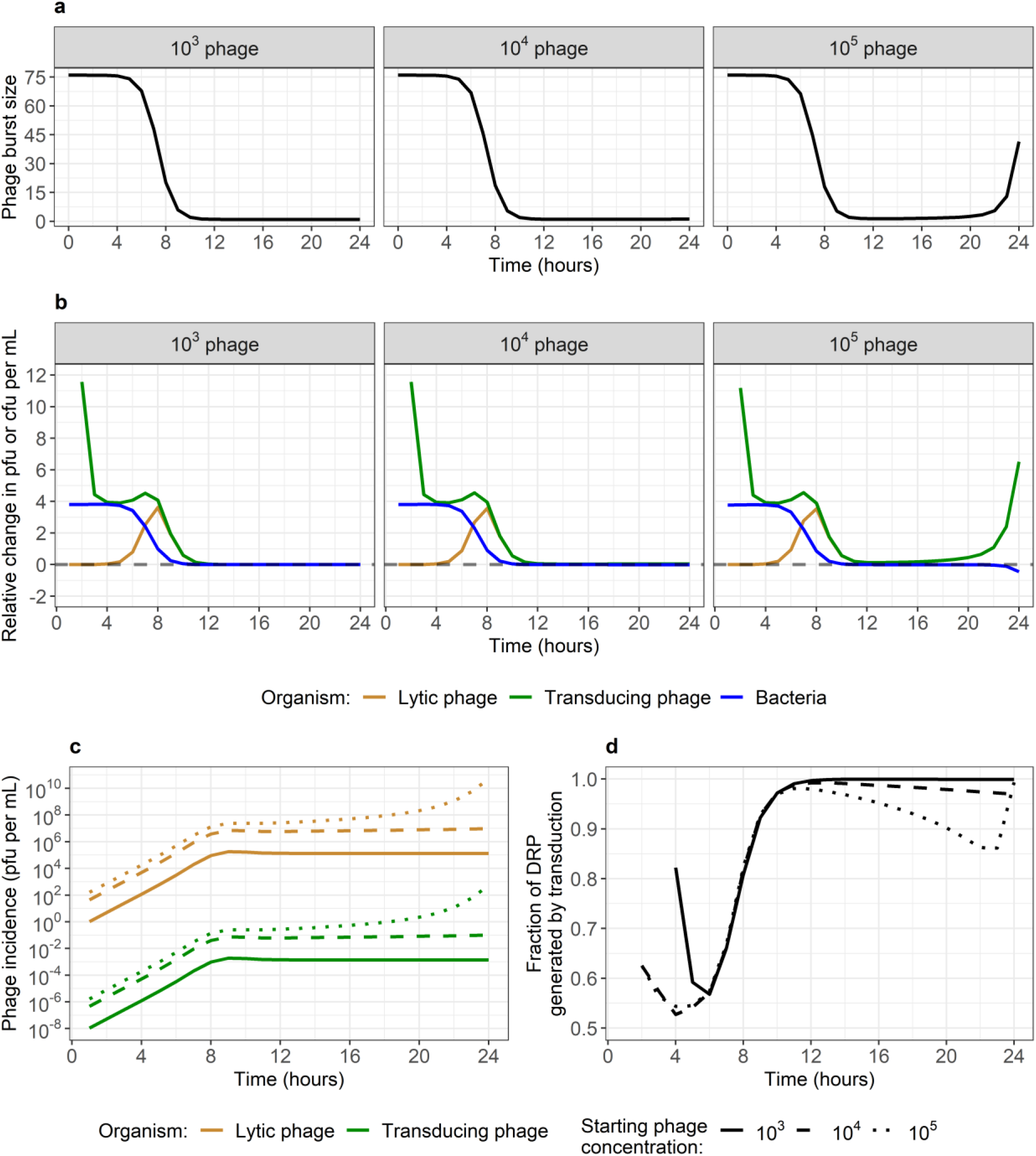
Underlying phage and bacteria dynamics generated by the best-fitting frequency-dependent model with burst size linked to bacterial growth. Model parameters are the median estimates from model fitting (Table 1). **(a) Phage burst size over time, by starting phage concentration.** As bacteria reach stationary phase after 8h, phage burst size decreases. In the 10^5^ dataset, we see that burst size is predicted to increase again after 20h. This is due to bacterial numbers decreasing as bacteria are being lysed by phage. **(b) Relative change in phage and bacteria numbers over time, by starting phage concentration.** The number of new phage generated at each time step increases (positive value) until bacteria reach stationary phase around 8h. This applies to lytic and transducing phage. In the 10^5^ dataset, phage keep increasing after 10h, eventually causing a decrease in bacterial numbers (negative value), which translates into a further acceleration in the increase in phage numbers due to the increased burst size (Figure 5a). After 8h, the relative changes in lytic and transducing phage numbers are identical. **(c) Incidence of lytic (gold) and transducing (green) phage over time, by starting phage concentration (linetype).** For any dataset and time-point, there is approximately 1 new transducing phage generated for each 10^8^ new lytic phage. **(d) Fraction of double-resistant progeny (DRP) generated by transduction each hour over time, by starting phage concentration (linetype).** DRP generation always occurs predominantly by transduction, rather than by growth of already existing DRP. Note that the time at which DRP are first generated varies by starting phage concentration.

We used our best-fitting model to reproduce our *in vitro* data (Figure 2) and uncover the underlying phage-bacteria dynamics. Due to the link between phage burst size and bacterial growth, burst size decreases as bacteria reach carrying capacity after 8h (Figure 6a-b). This is reflected in the relative change in phage numbers, which tends towards 0 after 8h (Figure 6b). After this point, phage incidence remains stable for the 10^3^ and 10^4^ pfu/mL dataset, but starts increasing again significantly after 20h for the 10^5^ pfu/mL dataset as bacteria numbers start decreasing due to phage predation, allowing burst size to increase again (Figure 6a-c).

We estimate that for every 10^8^ new lytic phage released during burst, there was approximately one transducing phage carrying an antibiotic resistance gene (Table 1, Figure 6c). Note that new double-resistant progeny (DRP) can either be generated by transduction, or by replication of already existing DRP. Using the model, we found that DRP were always predominantly generated by transduction rather than by growth (Figure 6d). This is because DRP only appear after 2 to 4h, while after 4h bacterial growth rate starts decreasing as the total bacteria population approaches carrying capacity (Figure 6b&d).

## Discussion

We observed rapid *in vitro* horizontal gene transfer of antimicrobial resistance (AMR) by generalised transduction in *Staphylococcus aureus*, alongside equilibriums in phage and bacteria numbers which varied depending on the starting number of phage. The most accurate mathematical model to replicate phage-bacteria dynamics, including generalised transduction, represented phage predation as a frequency-dependent interaction, and linked phage burst size to bacterial growth. To the best of our knowledge, these two elements have both been suggested previously ^17,18,34^, yet never combined.

Density-dependent models have been compared to data at less fine time scales (e.g. daily time points) or over smaller time periods (e.g. less than 8h), where they were able to reproduce *in vitro* values from experiments in chemostats, and have been helpful to improve our basic understanding of phage-bacteria dynamics ^14–16^. However, here we show that this type of interaction is not able to reproduce finer hourly dynamics, and does not perform consistently when varying concentrations of starting phage and bacteria. Using this, alongside a critique of the mathematical implications of this process, we argue that density-dependence is not a biologically accurate representation of phage predation, as it fails to reproduce these dynamics at high or low numbers of phage and bacteria, which would correspond to scenarios potentially seen during phage therapy.

Our work adds to the growing body of evidence that phage predation depends on bacterial growth ^14,17–23^. This has implications for antibiotic-phage combination therapy, as it suggests that bacteriostatic antibiotics, which prevent bacterial growth, could reduce phage predation. This effect has been previously seen in *S. aureus* ^38^. In the environment, including in persistent infections, bacteria spend most of their time in stationary phase ^39^. This suggests that bacteria and phage may be able to co-exist for prolonged periods of time in a broad range of settings, without the phage systematically eradicating the bacteria. Under such conditions, phage may mediate horizontal gene transfer by transduction between bacteria at relatively low levels, but for prolonged periods of time. This may be particularly relevant for *S. aureus*, since approximately 20% of humans are colonised asymptomatically by this bacterium at any given time ^40^, and at least 50% of these carriers may also carry phage capable of generalised transduction ^41^, suggesting a constant background evolution rate for *S. aureus* in the human population. Combined with environmental exposure to antibiotics which acts as a selective pressure, this may contribute to the risk of multidrug resistant bacteria evolution.

Our experimental design is both a strength and a limitation of our study. Since we jointly designed the experiments and models, we are confident that we have included in our mathematical model all the organisms and interactions present *in vitro*. We are therefore confident in the conclusions on model structure, however, the usage of such a specific experimental system with two bacterial strains of the same genetic background and one phage limits the generalisability of our parameter values, as these will likely vary for different bacteria and phage. Growth conditions will likely also differ between the *in vitro* environment studied here, and *in vivo* conditions. Here, our model assumes that phage do not decay, that bacteria do not become resistant to phage, and that they can grow indefinitely as they are observed in a rich medium for 24h only, but over longer periods of time it may be necessary to revisit these assumptions ^42^. Finally, we assumed that the proportion of transducing phage created was independent of the gene being transduced (*ermB*, on the bacterial chromosome, or *tetK*, on a plasmid). This was supported by preliminary work (not shown), but should be further investigated to improve our understanding of the factors that can facilitate or prevent transduction of different genes. To answer all of these questions, future work should investigate both phage predation and transduction dynamics over longer time periods, with different strains of bacteria and phage.

All our models captured certain aspects of the trends seen *in vitro*, but also underestimated phage numbers between 5-7h by up to 20 times. This is likely a consequence of our experimental design. To count lytic phage, we centrifuged and filtered the co-culture to remove bacteria. This could have caused the premature burst of some phage-infected bacteria, artificially increasing the numbers of phage we then counted ^43^. Since the period between 5-7h is when phage infections are highest (Figure 6b), this is why we would see such a large discrepancy at this stage. We also note that the models with only phage burst size linked to bacterial growth underestimated the number of double-resistant progeny (DRP). This small difference (up to 10 cfu/mL) is likely due to our choice of using a deterministic model. This type of model is useful for our purpose of fitting to *in vitro* data and analysing the underlying dynamics here, but mathematically allows for fractions of bacteria to exist, instead of just whole numbers. Future analyses using a stochastic model would better capture random effects, which can have an important impact at low numbers.

Multiplicity of infection (MOI, starting ratio of phage to bacteria) is a commonly used metric to present results of experiments with these organisms ^32^. With a starting concentration of 10^4^ bacteria per mL, we were able to fit our model to the dynamics for two MOI (0.1 and 10), and replicate those of a third (1). However, when trying to use the same model for these same three MOI, but with a starting bacterial concentration to 10^6^, we found differences between our model and values seen after 24h. This indicates that MOI is not appropriate to summarise all the complexity of the underlying phage-bacteria dynamics. Future experimental studies should express their results as a function of their starting concentration of phage and bacteria, not just MOI.

In any case, the failure of our model to replicate 24h values with a different starting bacteria concentrations shows that, whilst we have reduced the model structure uncertainty, we are still not fully capturing the phage-bacteria interaction. Currently, our model predicts that, for a starting concentration of 10^6^ bacteria, a starting concentration of 10^5^ phage or more will be enough to cause a decrease in bacterial numbers after 24h, while our data shows that the starting concentration of phage must be higher than 10^6^ for this to happen. *In vitro*, it is likely that slower bacterial growth simultaneously affects the phage adsorption rate, latent period and burst size, each to varying extents ^14,17–23^. This would explain why we would need a higher starting concentration of phage for a higher starting concentration of bacteria, to exert a strong enough predation pressure before bacteria reach stationary phase, causing a reduction in phage predation. However, here we have only made the first step in this process, having linked the burst size linearly to the bacterial growth rate, instead of trying to link different phage predation parameters to bacterial growth using different functions. These complexities need to be explored further, supported by i*n vitro* work measuring phage predation parameters at various time points. In *S. aureus*, wall teichoic acid (WTA) is the phage receptor ^44,45^. Lack of WTA glycosylation has been shown to induce phage resistance ^46^, and changes in WTA structure at different growth phases may be possible, since one of the genes involved in its synthesis is repressed by a quorum sensing system ^47^. However, to the best of our knowledge this has not yet been investigated.

Despite being recognised as a major mechanism of horizontal gene transfer, thus far there have been limited mathematical modelling studies on the dynamics of transduction of AMR ^13^. Using our model, we are able to estimate numbers of transducing phage which we cannot count *in vitro*, and see that approximately 1 generalised transducing phage is generated per 10^8^ lytic phage, consistent with previous estimates ^48,49^. Here, we show that this number, which may seem insignificant, is enough to consistently lead to the successful horizontal gene transfer of AMR, resulting in DRP after only 7h, substantially less than the usual duration of antibiotic treatment. We also show that transduction is the dominant mechanism to create new DRP throughout the entire experiment, rather than growth of existing DRP. This echoes the conclusions of previously published work on the importance of transduction, including *in vivo* experiments and with other *Staphylococcus* species ^6,7,29,50^.

Our findings suggest that transduction is currently under-emphasised in the exploration of phage-bacteria dynamics. Future studies on this topic should not assume that transduction can be dismissed by default, but instead investigate whether it is relevant in their system. This requires further *in vitro* and *in vivo* monitoring to identify scenarios where transduction plays a significant role in the transfer of AMR genes, likely depending on the environment, and characteristics of the bacteria and phage present. This will require new experimental designs, since counting phage numbers can be difficult, notably with clinical strains of bacteria. This should also be investigated in the presence of antibiotics, where the importance of selection enters, increasing the fitness of the small numbers of DRP generated by transduction.

In conclusion, the dual nature of phage (predation and transduction) leads to complex interactions with bacteria. These dynamics must be clarified, to correctly evaluate the extent to which phage contribute to the global spread of AMR. We must also understand this dual nature to guarantee a safe design of phage therapy. Otherwise, ignoring transduction may lead to worse health outcomes in patients if phage contribute to spreading AMR instead of overcoming it. Interdisciplinary work will be essential to answer these urgent public health questions in the near future.

## Materials and Methods

All analyses were conducted using the statistical software R ^51^. The underlying code and data are available in a GitHub repository: https://github.com/qleclerc/mrsa_phage_dynamics.

### Experimental methods

#### Strains and phage used

The *Staphylococcus aureus* parent strains used for our transduction experiment were obtained from the Nebraska Transposon Mutant Library ^52^. These were strain NE327, carrying the *ermB* gene encoding erythromycin resistance and knocking out the ϕ3 integrase gene, and strain NE201KT7, a modified NE201 strain with a kanamycin resistance cassette instead of the *ermB* gene knocking out the ϕ2 integrase gene, and a pT181 plasmid carrying the *tetK* gene encoding tetracycline resistance ^53^. Growing these strains together in identical conditions as our co-culture below, but without the addition of exogenous phage, does not lead to detectable horizontal gene transfer (HGT; data not shown). To enable HGT, exogenous 80α phage was used, a well-characterised temperate phage of *S. aureus* capable of generalised transduction ^33^. To count lytic phage, *S. aureus* strain RN4220 was used, a restriction deficient strain highly susceptible to phage infection ^54^.

#### Transduction co-culture protocol

Pre-cultures of NE327 and NE201KT7 were prepared separately in 50mL conical tubes with 10mL of Brain Heart Infusion Broth (BHIB, Sigma, UK), and incubated overnight in a shaking water bath (37°C, 90rpm). The optical densities of the pre-cultures were checked at 625nm the next day to confirm growth. The pre-cultures were diluted in phosphate-buffered saline (PBS), and added to a glass bottle of fresh BHIB to reach the desired starting concentration in colony forming units per mL (cfu/mL) for each strain, forming a master mix for the co-culture. CaCl2 was added at a concentration of 10mM to the master mix. Phage 80α stock was diluted in phage buffer (50 mM Tris-HCl pH 7.8, 1 mM MgSO4, 4 mM CaCl2 and 1 g/L gelatin; Sigma–Aldrich), and added to the master mix to reach the desired starting concentration in plaque forming units per mL (pfu/mL). Ten 50mL conical tubes were prepared (one co-culture tube for each timepoint, from 0 to 8h and 16 to 24h), each with 10mL from the master mix. Each co-culture tube was then incubated in a shaking water bath (37°C, 90rpm) for the corresponding duration.

Bacteria counts for each timepoint were obtained by diluting the co-culture in PBS before plating 50μL on selective agar, either plain Brain Heart Infusion Agar (BHIA, Sigma, UK), BHIA with erythromycin (Sigma, UK) at 10mg/L, BHIA with tetracycline (Sigma, UK) at 5mg/L, or BHIA with both erythromycin and tetracycline (10mg/L and 5mg/L respectfully). Note we plated 500μL instead of 50 on the plates with both antibiotics, to increase the sensitivity of the assay. This allowed distinction between each parent strain, resistant to either erythromycin or tetracycline, and the double resistant progeny (DRP) generated by transduction. Plates were then incubated at 37°C for 24h, or 48h for plates containing both antibiotics. Colonies were counted on the plates to derive the cfu/mL in the co-culture for that timepoint.

Lytic phage counts for each timepoint were obtained using the agar overlay technique ^55^. Briefly, the co-culture was centrifuged at 4000rpm for 15 minutes, filtered twice with 10μm filters, and diluted in Nutrient Broth No. 2 (NB2, ThermoFisher Scientific, UK). 15mL conical tubes were prepared with 300μl of RN4220 grown overnight in NB2, and CaCl2 at a concentration of 10mM. 200μl of diluted phage were added, and the tubes were left to rest on the bench for 30 minutes. The contents of the tubes were then mixed with 7mL of phage top agar, and poured on phage agar plates. Phage agar was prepared using NB2, supplemented with agar (Sigma, UK) at 3.5g/L for top agar and 7g/L for plates. The plates were incubated overnight at 37°C. Clear spots in the bacterial lawn were counted to derive the pfu/mL in the co-culture for that timepoint.

#### Polymerase chain reaction protocols

To confirm that DRP contained both the *ermB* and *tetK* genes, primers ermBF (5′- CGTAACTGCCATTGAAATAGACC-3′), ermBR (5′-AGCAAACTCGTATTCCACGA-3′), tetKF (5′-ATCTGCTGCATTCCCTTCAC-3′), and tetKR (5′-GCAAACTCATTCCAGAAGCA-3′) were used. Strains NE327 (only containing *ermB*) and NE201KT7 (only containing *tetK*) were used as positive and negative controls.

To confirm that 80α lysogeny did not occur in our co-culture, we applied a previously published method ^33^ and used a combination of four primers: SaRpmF (5′-GACTGAATGCCCAAACTGTG-3′) in the *S. aureus rpmF* gene, SMT178 (5’-GGCTGGGAATTAATGGAAGATG-3′) in the 80α integrase, SaSirH (5′- TTAAGTAGCATCGTTGCATTCG-3′) in the *S. aureus sirH* gene, and SMT179 (5′- GAGTCCTGTTTGCGAATTAGG-3′) in the 80α ORF73 region. SaRpmF and SMT178 were used to amplify the left prophage junction (*attL*), SaSirH and SMT179 to amplify the right junction (*attR*), and SaRpmF and SaSirH to amplify the bacterial insertion site (*attB*). RN4220 was used as a negative control for lysogeny, and JP8488, an RN4220 strain lysogenic for 80α, was used as a positive control (obtained from José Penadés and Nuria Quiles, Imperial College London).

All PCRs were conducted using OneTaq Hot Start Quick-Load 2X Master-mix, following the manufacturer’s protocol. Tested samples were homogenised in 20μl nuclease-free water, and 1.5μl of each suspension was used as template for a total reaction volume of 25μl.

### Mathematical modelling methods

#### General model structure

We designed a deterministic, compartmental model to replicate our experimental conditions. We included 6 populations: B_E_ (corresponding to ery-resistant NE327), B_T_ (tet-resistant NE201KT7), B_ET_ (double resistant progeny, DRP), P_L_ (lytic phage), P_E_ (phage transducing *ermB*) and P_T_ (phage transducing *tetK*). Their interactions are represented in Figure 2.

Bacteria from each strain θ (θ ∈ {E, T, ET}) can multiply at each time step *t* following logistic growth at rate μ_θ_, with a maximum value μ_maxθ_ which declines as the total bacteria population N (= B_E_ + B_T_ + B_ET_) approaches carrying capacity N_max_.

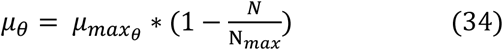

At each time step *t*, a proportion λ of lytic phage (P_L_) infect a number of bacteria (φ_L_), replicate, and burst out from the bacteria with a burst size δ + 1 after a latent period τ. During phage replication, a proportion α of new phage are transducing phage. The nature of the transducing phage (P_E_ or P_T_) depends on the bacteria being infected (e.g. B_E_ bacteria can only lead to P_E_ phage). Then, a proportion λ of these transducing phage (P_E_ or P_T_) infect a number of bacteria (φ_E_ or φ_T_). If they successfully infect a bacterium carrying the other resistance gene (e.g. P_E_ phage infecting a B_T_ bacterium), this creates double resistant progeny (B_ET_). The complete model equations can be found below.

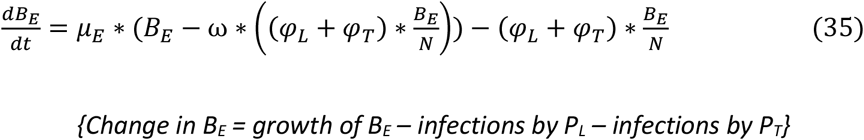

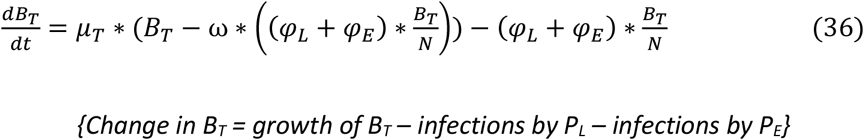

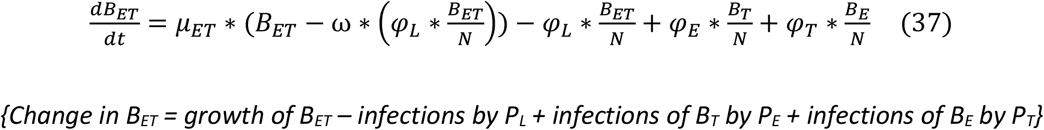

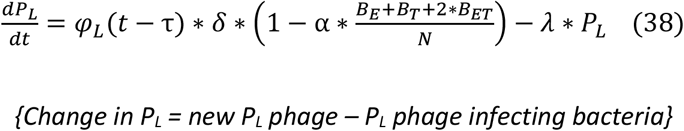

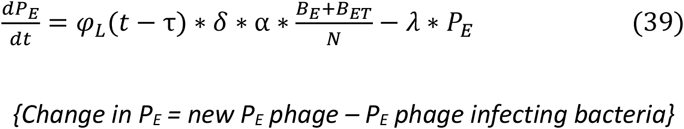

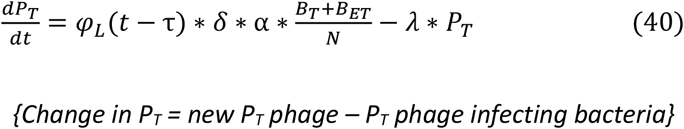

Some parameters (τ, α, ω) are constant, while others (μ_E_, μ_T_, μ_ET_, φ, β_L_, φ_E_, φ_T_, δ) can change at each time step and depending on the specified interaction mechanism. Note that ω is a special parameter equal to 0 if the model is density-dependent, or 1 if it is frequency-dependent.

#### Density-dependent interaction

Over one time step, both the number of phage infecting bacteria and the number of bacteria infected by phage are equal to the product of the number of phage, bacteria, and phage adsorption rate. In our equations for density-dependence, given the phage adsorption rate β, the proportion λ of phage that infect any bacteria is:

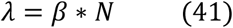

And the number of bacteria infected by a phage θ(θ ∈ {L, E, T}) is:

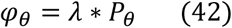

Note that the parameter ω is set to 0 in this case.

#### Frequency-dependent interaction

Using this interaction prevents the number of phage infecting bacteria over one time step to be higher than the total number of phage in the system (and the number of bacteria being infected one time step to be higher than the total number of bacteria in the system). Equations (41) and (42) then become:

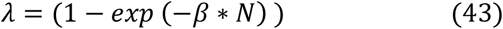

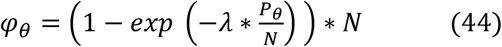

With the frequency-dependent interaction, we set the parameter ω to 1. This ensures that, over one time step and for any bacterium, phage infection and bacteria replication are mutually exclusive events. Without this modification, phage infections would not be able to reduce bacterial population size due to mathematical constraints. Equation (45) shows a simplified frequency-dependent model for a single bacterial strain *B*, without any correction term.

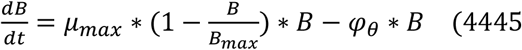

According to equation (45), the change in bacterial numbers depends on the relative values of bacterial growth and bacterial death due to phage predation, expressed in equation (46).

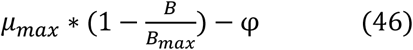

The necessity for a correction term on the left hand side of equation (45) arises from the maximum values of μ_max_ and φ_θ_. As can be deduced from equation (44), the maximum possible value for φ_θ_ is 1. According to our fitted parameter values, the maximum value for μ_max_ in our model is approximately 1.5, and the carrying capacity Bmax is approximately 2.8 × 10^9^. Substituting these into equation (46) leads to equation (47).

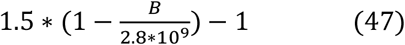

To have a decline in the bacterial population, we must therefore satisfy the condition stated in equation (48).

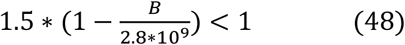

This can be rearranged into equation (49) by dividing by 1.5.

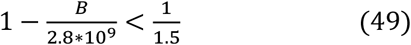

Finally, by subtracting 1 and multiplying by −2.8 × 10^9^, we obtain equation (50).

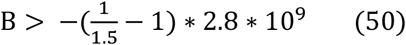

Solving equation (50) leads to the condition that the number of bacteria must be greater than 9.3 × 10^8^ in order for bacterial growth to be low enough that bacteria numbers decrease. In other words, it is impossible for the effect of phage predation to decrease bacteria numbers below 9.3 × 10^8^. We overcome this by applying a correction term to equation S1, leading to equation (51).

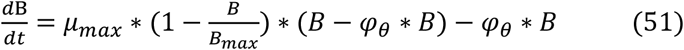

This is equivalent to saying that bacteria infected by phage cannot replicate and hence is more biologically realistic.

#### Link between bacterial growth and phage predation

We consider that reduced bacterial growth can lead to decreased phage predation, through reduced adsorption (β) and/or burst size (δ). Equations (52) and (53) allow these parameters to decrease as bacterial growth decreases, using the same principle of logistic growth as seen in equation (34).

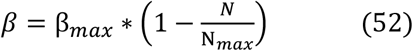

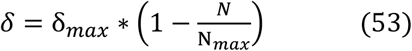

If we do not link these parameters to bacterial growth, we assign them their maximum values.

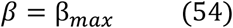

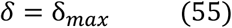

#### Model fitting

We fit our model to the *in vitro* data using the Markov chain Monte Carlo Metropolis–Hastings algorithm. For every iteration, this algorithm slightly changes the parameter values, runs the model, assesses the resulting model fit to the data, and accepts or rejects these new parameter values based on whether the model fit is better or worse than with the previous set of values. We run the algorithm with two chains, and once convergence has been reached (determined using the Gelman-Rubin diagnostic, once the multivariate potential scale reduction factor is less than 1.2 ^56^), we generate 50,000 samples from the posterior distributions for each parameter.

In a first instance, we used our growth co-culture data, where phage are absent, to calibrate the bacterial growth rate parameters μ_maxθ_ for each bacteria strain θ (θ ∈ {E, T, ET}), as well as the carrying capacity N_max_ using a simple logistic growth model (equation (9)). All other parameters related to phage predation were set to 0.

The phage predation parameters (τ, α, β_max_, δ_max_) were jointly estimated by fitting to the phage and double resistant bacteria numbers from the transduction co-culture data. Fitting was performed by evaluating the log-likelihood of each *in vitro* data point being observed in a Poisson distribution, with the corresponding model data point as a mean.

To mirror our experimental sampling variation, *in vitro* data points were scaled down to be between 1 and 100 before fitting, with the same correction applied to the corresponding model-predicted value for the same timepoint. For example, if at 1h there are 1.4 × 10^4^ phage *in vitro*, this is scaled down to 14, and if the corresponding model value is 5.3 × 10^6^, this is scaled down by the same magnitude (i.e. 10^3^), resulting in a value of 5300.

Previous research estimated that the latent period for 80α in *S. aureus* was approximately 40mins (0.67h), and that the burst size was approximately 40 phage per bacterium ^37^. Since this study did not provide error values for these point estimates, we assumed the standard deviation and chose the following informative priors for these parameters: τ ~ *Normal(0.67, 0.07)* (95% confidence interval: 0.53-0.81) and δ_max_ ~ *Normal(40, 7)* (95% confidence interval: 54-26). Due to a lack of available data, we used uninformative priors for the remaining parameters: α ~ *Uniform(0, 1)* and β_max_ ~ *Uniform(0, 1)*.

## Acknowledgments

Q.J.L and J.W. were supported by a studentship from the Medical Research Council Intercollegiate Doctoral Training Program (MR/N013638/1). A.G. and G.M.K were supported by grants from the Medical Research Council (MR/P028322/1 and MR/P014658/1 respectively). We would like to thank José Penadés and Nuria Quiles for providing the JP8488 *S. aureus* strain.

