## Supplementary Material for "The dual nature of bacteriophage: growth-dependent predation and generalised transduction of antimicrobial resistance"

### Supplementary Figures

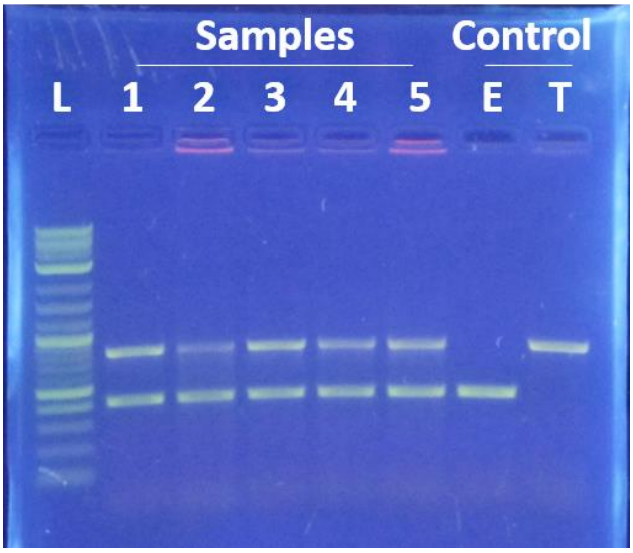


**Figure 1 – figure supplement 1: Confirmation of double-resistant progeny by polymerase-chain reaction.** Five single colonies were sampled from a double antibiotic plate (1-5), containing bacteria plated after 24h of co-culture started only with single-resistant parent strains (E and T) and exogenous phage. L: ladder; E: erythromycin resistance gene (*ermB*); T: tetracycline resistance gene (*tetK*).

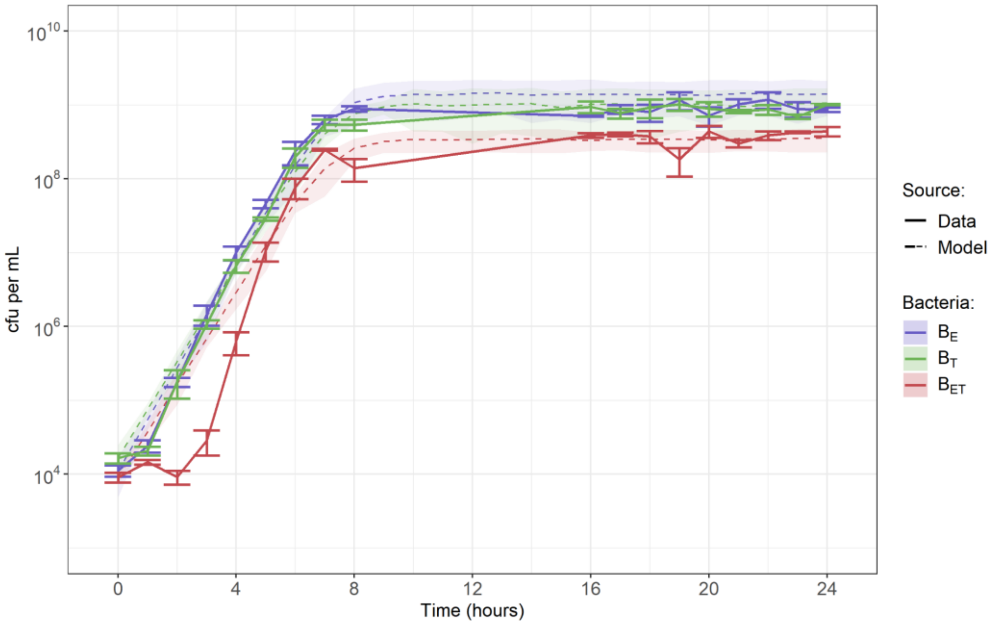


**Figure 1 – figure supplement 2: Growth curves for bacteria in the absence of exogenous phage.** $B_E$: bacteria resistant to erythromycin, $B_T$: bacteria resistant to tetracycline, $B_{ET}$: bacteria resistant to both erythromycin and tetracycline. Solid lines correspond to *in vitro* data, and dashed lines to the model output generated using the median values of the parameter distributions obtained by model fitting. Shaded areas indicate error obtained by resampling the model results from a Poisson distribution ten times.

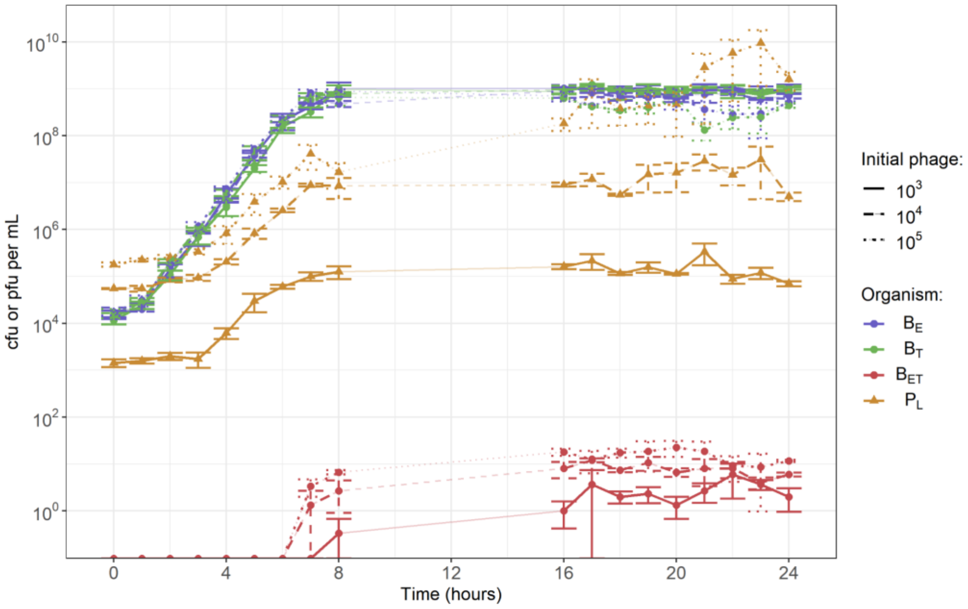


**Figure 1 – figure supplement 3: Transduction co-culture datasets overlaid.** The starting concentration of both single-resistant *S. aureus* parent strains ($B_E$ to erythromycin & $B_T$ to tetracycline) is $10^4$ colony-forming units (cfu) per mL. The starting concentration of exogenous phage 80$\alpha$ ($P_L$) is either $10^3$ (solid lines), $10^4$ (dashed) or $10^5$ (dotted) plaque-forming units (pfu) per mL. Error bars indicate mean +/- standard error, from 3 experimental replicates. There is no data for the time period 9h-15h.

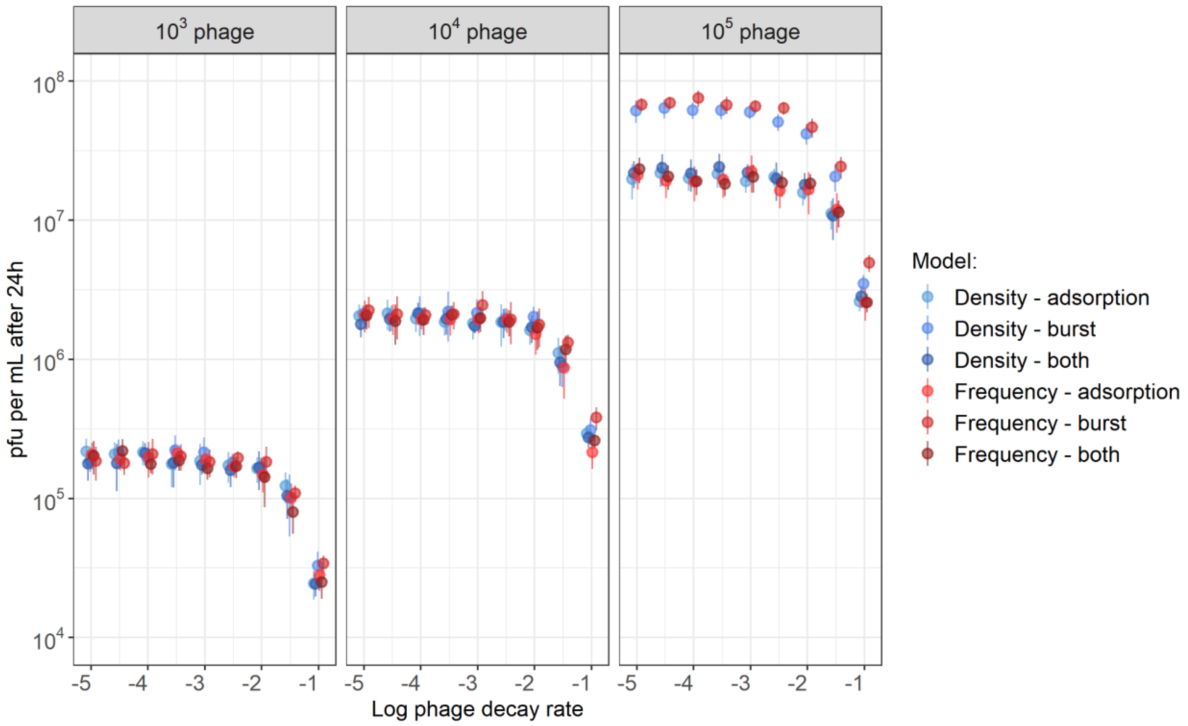


**Figure 5 – figure supplement 1: Model results are not affected by phage decay rate over a wide range of values.** Previous estimates of phage decay rate per hour are between $10^{-3}$ *in vitro* and up to $10^{-1}$ *in vivo* [38]. Models are either density or frequency-dependent, with either or both the phage adsorption rate and burst size linked to bacterial growth.

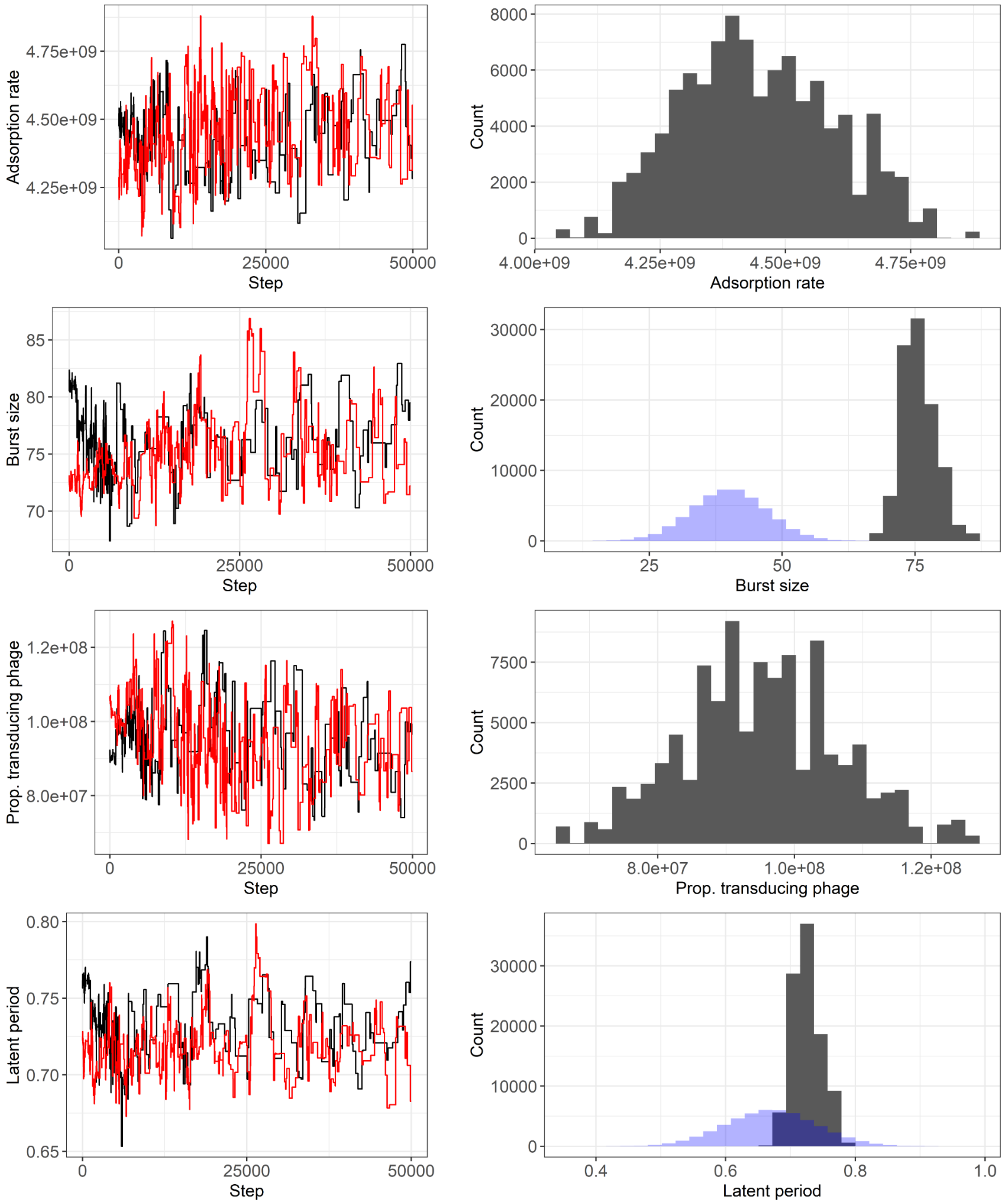


**Figure 5 – figure supplement 2: Convergence and posterior distributions for the best-fitting model (with a frequency-dependent interaction and a link between phage burst size and bacterial growth). (a-d) Parameter convergence plots.** Fitting was performed using two chains (black and red). **(e-h) Posterior distributions.** The prior distributions for the burst size and latent period are shown in blue. The prior distributions for other parameters were uninformative and not shown.

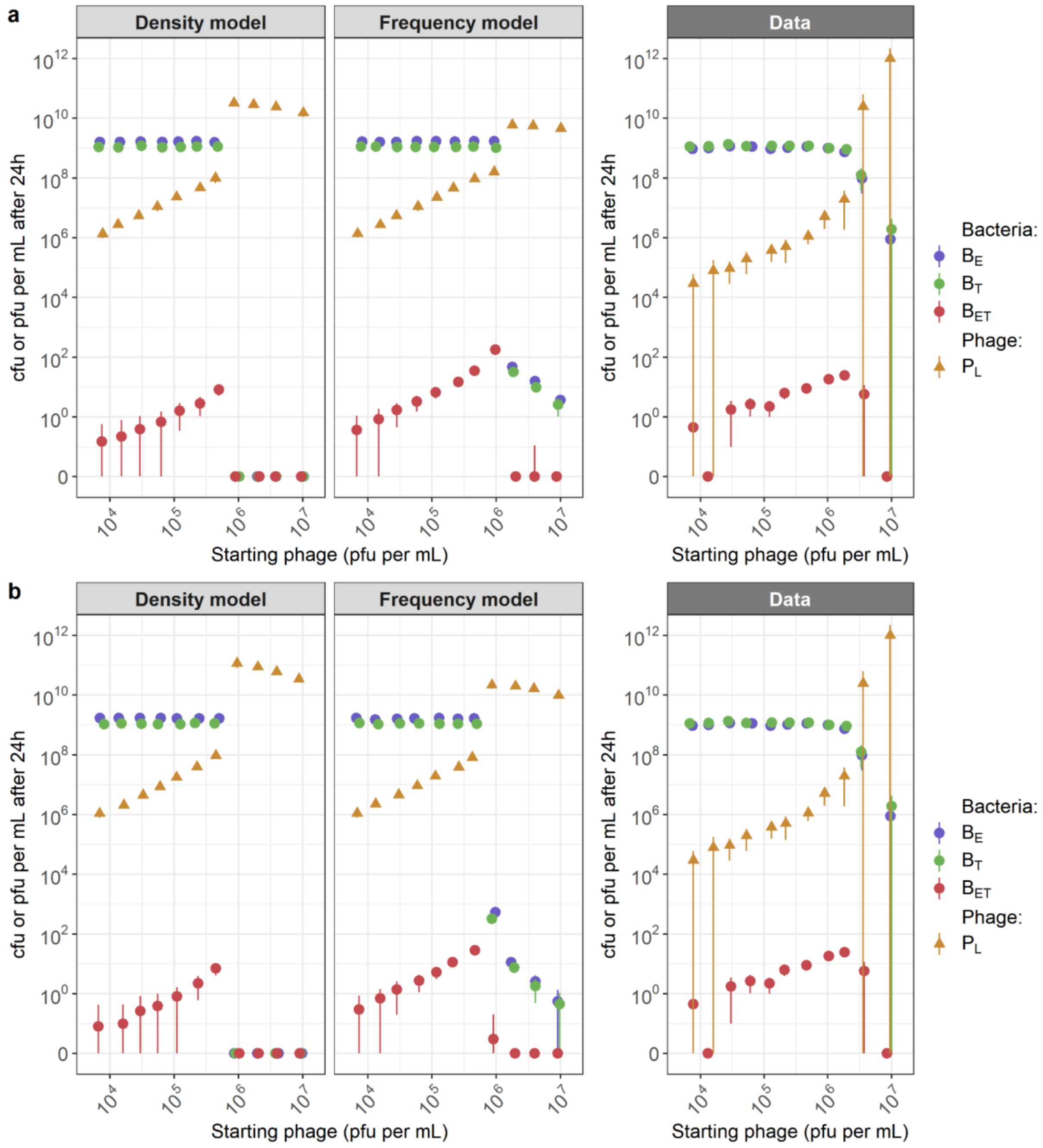


**Figure 5 – figure supplement 3: Model performance in reproducing the 24h data values for different starting concentrations of phage with different links between phage predation and bacterial growth rate: A) Phage adsorption rate decreases as bacterial growth rate decreases. B) Phage burst size and adsorption rate decrease as bacterial growth rate decreases.** Phage predation is either density- or frequency-dependent in the models. Model parameters are those estimated for the corresponding model as shown in Table 1. In the co-culture used to generate the data, each single-resistant parent strain ($B_E$ and $B_T$) is added at a starting concentration of $10^6$ cfu/mL, and no double-resistant progeny ($B_{ET}$) are initially present. The starting concentration of lytic phage ($P_L$) varies (x axis).
